# The cyanobacterial saxitoxin exacerbates neural cell death and brain malformations induced by Zika virus

**DOI:** 10.1101/755066

**Authors:** Carolina da S. G. Pedrosa, Leticia R. Q. Souza, Caroline V. F. de Lima, Pitia F. Ledur, Karina Karmirian, Tiago A. Gomes, Jimena Barbeito-Andres, Marcelo do N. Costa, Luiza M. Higa, Maria Bellio, Flavio A. Lara, Amilcar Tanuri, Patricia P. Garcez, Arnaldo Prata-Barbosa, Fernanda Tovar-Moll, Renato J. R. Molica, Stevens K. Rehen

## Abstract

The northeast (NE) region of Brazil commonly goes through drought periods, which favor cyanobacterial blooms, capable of producing neurotoxins with implications for human and animal health. The most severe dry spell in the history of Brazil occurred between 2012 and 2016. Coincidently, the highest incidence of microcephaly associated with the Zika virus (ZIKV) outbreak was described in the NE region of Brazil during the same years. In this work, we tested the hypothesis that saxitoxin (STX), a neurotoxin produced in South America by the freshwater cyanobacteria *Raphidiopsis raciborskii*, could have contributed to the most severe Congenital Zika Syndrome (CZS) profile described worldwide. Quality surveillance showed higher cyanobacteria amounts and STX occurrence in human drinking water supplies of NE compared to other regions of Brazil. Experimentally, we described that STX doubled the amount of ZIKV-induced neural cell death in progenitor areas of human brain organoids, while the chronic ingestion of water contaminated with STX before and during gestation caused brain abnormalities in offspring of ZIKV-infected immunocompetent C57BL/6J mice. Our data indicate that saxitoxin-producing cyanobacteria is overspread in water reservoirs of the NE and might have acted as a co-insult to ZIKV infection in Brazil. These results raise a public health concern regarding the consequences of arbovirus outbreaks happening in areas with droughts and/or frequent freshwater cyanobacterial blooms.

**Author summary:** The uncontrolled spreading of cyanobacteria in drinking water reservoirs has been the cause of serious public health problems worldwide. Toxin-producing cyanobacterial blooms commonly occur during drought periods in the northeast (NE) region of Brazil. During Zika Virus (ZIKV) outbreak in 2015-16, Brazilian NE showed disproportionately higher microcephaly incidence. Here, we test the hypothesis that the cyanotoxin saxitoxin (STX) may act as a co-insult for ZIKV. Water quality surveillance data showed increased cyanobacteria population and higher STX amount in NE region during 2014-2018. *In vitro*, we observed that neural progenitor cell death was doubled after STX exposure to ZIKV-infected brain organoids. *In vivo*, chronic ingestion of STX during gestational period potentiated ZIKV-derived brain abnormalities in newborn mice. Our study provides new insights that may explain the discrepancies among Brazilian regions regarding CZS severity. Moreover, the data highlight the importance of cyanobacteria and cyanotoxin freshwater monitoring for future arbovirus outbreaks.

## Introduction

Human population growth, associated with disorderly occupation of territory, results in waste discard in the freshwater reservoir. This Ambiental problem could be escalated by long periods of drought, leads to aquatic ecosystems eutrophication, with the main problem being the mass proliferation of cyanobacteria (blooms) (1). Cyanobacterial blooms comprise hepatotoxin- and neurotoxin-producing species responsible for wild and domestic animals intoxication, besides the contamination of human drinking water supplies (2). Previous studies have shown that 60% of all fresh water samples containing cyanobacteria used to be toxic, with neurotoxin-producing ones being more common in North America, Europe and Australia (3).

Brazilian northeast (NE) usually faces periods of severe drought, with the most severe ever recorded occurring between 2012 and 2016 (4). Besides reducing the reservoirs to critical volumes, which results in water supply deficiency (5), this rainy scarcity favors cyanobacterial blooms (6, 7). A literature survey of cyanobacteria publication from 1930 to 2016 showed that the highest number of toxic bloom events occurred in Pernambuco (PE) state, where was described the presence of microcystins, cylindrospermopsin, five variants of saxitoxin (STX) and anatoxin-a(S) in freshwater (8). Extreme climate events promote changes in the dominance of cyanobacteria (7) as shown during the drought in 1998 (a consequence of the El Niño in 1997), which favored the proliferation of *Raphidiopsis raciborskii* (formely *Cylindrospermopsis raciborskii*) (9) in almost 40 reservoirs in the NE of Brazil (6). The *R. raciborskii* has high adaptability to unfavorable conditions because of its physiological characteristics, that includes akinete formation and tolerance to low phosphorus and nitrogen availability (10). Most important, saxitoxin producing strains of *R. raciborskii* where positively selected among non-producing strains in NE superficial freshwater reservoir, as STXs would serve as a protection against water high salinity and or hardness (11–13).

The Brazilian strain of *R. raciborskii* produces STX, one of the most potent paralytic shellfish toxin (PST) found in freshwater and marine ecosystems (10). PSTs are a group of neurotoxic alkaloids that act binding to voltage-gated sodium channels, blocking the generation of action potentials in neurons. The acute exposition to high amounts of PST results in numbness and even death by respiratory failure (14). In contrast, little is known about the effects of chronic low-dose exposure to PSTs (15). Because of their aforementioned dangerousness, a safety level of 3 μg/L of STX has been established in Brazilian water quality guidelines (16). However, *in vitro* exposure to low levels of STX has already been reported to result in impaired neurite outgrowth and altered expression of proteins related to cell apoptosis and mitochondrial function (15,17).

The amount of STX usually found in reservoirs of the Brazilian semi-arid region varied between 0.003 and 0.766 μg/L, depending on the period of the year (18). In 2000, during a toxic bloom at the northeast state of Rio Grande do Norte, *R. raciborskii* represented 90-100% of total phytoplankton species (19). In case of severe water scarcity, the most impoverished population uses raw water from alternative sources without effective elimination of microorganisms. The consumption of water from ponds, water trucks, wells and household water reservoirs has already been associated with diarrhea outbreaks in states of the Brazilian NE (20). Furthermore, it is important to notice that STX could also accumulate in marine organisms such as freshwater fishes, which is the main animal protein source of many NE communities (21). The effects of this accumulation in humans are not completely understood.

Zika virus (ZIKV) infection became an international concern when it was linked to a high rate of congenital brain abnormalities in Brazil (22,23). The incidence of microcephaly varied among regions, with the highest frequency being found in the NE of Brazil (24,25) (S1 Fig in Supporting Information). In contrast, the total number of cases of ZIKV infection was lower in NE compared to middle-west or southeast (SE) regions (26). Authors have suggested that a co-factor could be acting with ZIKV, contributing to this divergence among NE and other regions of Brazil; however, none has been confirmed until now (27).

The present study aimed to evaluate cyanobacteria and STX spreading among Brazilian regions during the ZIKV outbreak; confirming the synergism between STX and ZIKV *in vitro* using human brain organoids, and *in vivo*, using low-dose exposition of STX to mice that may exacerbate the neurological consequences of viral congenital infection. Our results show that STX occurred in almost half of municipalities in the Brazilian NE, while the majority of other regions presented STX in less than 5%. STX combined with ZIKV increased neural cell death and brain malformations, *in vitro* and *in vivo*. Therefore, STX could be an environmental co-factor associated with the highest incidence of brain abnormalities caused by ZIKV in the northeast of Brazil compared to any other region of the world.

## Methods

### Occurrence of cyanobacteria and STX in water reservoirs of Brazil

The data about the number of cyanobacteria and STX presence were obtained from SisAgua - Water Quality Surveillance Information System for Human Consumption, a Brazilian Ministry of Health integrated data bank. The number of cyanobacteria per milliliter was determined in water reservoir destinated to human use, before treatment, from 2014 to 2018. Values were compiled and corrected by the number of municipalities in each Brazilian state. Then, the percentage of the measurements below 10,000 cells/mL, between 10,000 and 20.000 cells/mL and above 20,000 cells/mL per municipality were organized per each region of Brazil. STX presence at treated water from 2014 to 2018 was compiled the same way as cyanobacteria concentration and their presence per municipality were organized per each region of Brazil.

### ZIKV propagation and titration

ZIKV (Recife/Brazil, ZIKV PE/243, number: KX197192.1) was provided by Dr. Marli Tenório Cordeiro from Fundação Oswaldo Cruz/Centro de Pesquisas Aggeu Magalhães, Brasil. The procedure of ZIKV isolation was described previously (28). The virus was propagated in C6/36 *Aedes albopictus* cell line at multiplicity of infection (MOI) of 0.01 and cultured for 6 days in Leibovitz’s L-15 medium (Thermo Fisher Scientific, Waltham, MA) supplemented with 0.3% tryptose phosphate broth (Sigma-Aldrich), 2 mM glutamine and 1x MEM non-essential amino acids (Thermo Fisher Scientific) and 2% FBS. ZIKV titers were determined by conventional plaque assay.

### Human brain organoids

Human induced pluripotent stem (iPS) cells were obtained from Coriell Institute for Medical Research repository (GM79A). iPS cells were cultured in mTeSR1 media (StemCell Technologies, Vancouver, CAN) on the top of Matrigel (BD Biosciences, Franklin Lakes, NJ). When colonies reached 70-80% confluence, iPS cells were dissociated with Accutase (MP Biomedicals, Santa Ana, CA), centrifuged at 300g, resuspended in media and counted. 9,000 cells/well were plated in ultra-low attachment 96-well plates and maintained at 37°C and 5% CO_2_.

Next day, medium was replaced with hESC media and the embryoid bodies (EBs) were cultured for 6 days as previously described (29). Then, EBs were transferred to 24-well ultra-low attachment culture plates containing Neural Induction Media: 1% N2 Supplement (Thermo Fisher Scientific), 1% GlutaMAX (Life Technologies), 1% P/S, heparin 1 μg/mL for 4 days. Organoids were coated with Matrigel during 1 hour at 37 °C and 5% CO_2_ and returned to 24-well ultra-low attachment plates in Neurodifferentiation Media (NDM) with no vitamin A for additional 4 days in static culture and subsequently, transferred to agitation in NDM with vitamin A until day 50. Culture media changes were performed weekly.

### ZIKV infection in human brain organoids

The superficial cell number in organoids was calculated by dividing the superficial area (calculated using: 4πr2) by the mean cell area in the organoid surface (15 μm^2^). Brain organoids were infected using ZIKV MOI 0.5 (2−6.5 × 10^5^ PFU per organoid) - for 2 h, then cultured in medium with (or without) STX 12 μg/L (NRC Halifax, CAN) for 13 days. Mock-exposed organoids (treated and non-treated with STX) were used as control. The assay was performed in triplicates in three independent experiments.

### Animal experimental design, STX exposition and ZIKV infection

C57BL/6J ZIKV-refractory (30,31) nulliparous female 6-week-old mice received standard filtered water *ad libitum* supplemented (or not) with 15 ng/L of STX 7-10 days before mating and until harvesting date. All females were fed a standard diet with the recommended amount of macro and micro-nutrients (TD91352, Harlan Teklad, Madison). No significant differences in water intake were observed between groups.

Pregnancy was confirmed through observation of post-coital vaginal plug for estimation of embryonic age. ZIKV (virus plus C6/36 cell line supernatant) or Mock (C6/36 cell line supernatant) was administered intraperitoneally on E12. ZIKV groups received 10^6^ plaque-forming units per animal. Harvesting of samples was carried out on the first day of postnatal life (P0).

### Sample preparation for optical microscopy

Brain organoids and newborn mice brains were fixed in 4% paraformaldehyde solution (Sigma-Aldrich) for 2h and 48h, respectively. Organoids were cryopreserved in sucrose solution, immersed in O.C.T compound (Sakura Finetek, Netherland) and frozen at −80 °C, being sectioned at 20 μm slices in a Leica CM1860 cryostat for analysis. Newborn brains were embedded in 5% agarose/PBS (Bioline, Taunton, MA), being sectioned coronally at 80 μm in a vibrating microtome (VT1000S, Leica, Germany) for analysis.

### Immunofluorescence staining

After washing with PBS, sections were incubated in permeabilization/blocking solution (0.3% Triton X-100/ 3% goat serum, for organoids; 0.2% Triton X-100/ 2% goat serum, for newborn brains) for 2 h. Primary antibodies Mouse IgG anti-NS1 (1:500 - organoids or 1:10 – newborn brains; BF-1225-36 – BioFront Technologies, Tallahassee, FL) combined with rabbit IgG anti-Nestin (1:1000; RA22125 – Neuromics, Minneapolis, MN) in organoids or with, rabbit IgG anti-cleaved caspase 3 (1:300; 9661S - Cell Signaling, Danvers, MA) in newborn brains were incubated at 4°C overnight. Then, sections were incubated with secondary antibody goat anti-rabbit AlexaFluor 488 (1:400; A-11008 - Thermo Fischer Scientific), for organoids and newborn brains; and goat anti-mouse AlexaFluor 546 (1:500; A-11003 - Life technologies), for newborn brains, for 2 h. Nuclei were stained with 0.5 μg/mL 4’-6-diamino-2-phenylindole (DAPI) for 20 min. Apoptotic cells were stained with ApopTag® Red in Situ (S7165, Merck Millipore) according to manufacturer’s instructions.

10-12 periventricular nestin-positive fields of 100 μm^2^ of three sections per organoid from each experimental group were analyzed. Images were acquired on a confocal microscope Leica TCS SP8. The number of TUNEL positive cells per area was quantified using Cell Profiler Software (BROAD Institute, Cambridge, MA). Two sections per animal brain from each experimental group (three brains per group) were analyzed. Images were taken on microscope AxioImager A.1 (Zeiss, Oberkochen, DEU). Image J was used to determine the thickness of cortical layers.

### Ethical statements

Animals were housed in the Animal Care Facility of the Microbiology Institute of the Federal University of Rio de Janeiro. Protocols for animal handling were approved by the Research Ethics Committee of the Federal University of Rio de Janeiro (CONCEA registration number 01200.001568/2013-87, acceptance number 037/16).

### Statistical analysis

*In vitro* and *in vivo* results were expressed as mean plus standard error of the mean (SEM). Data sets were compared using One-Way ANOVA, followed by post-test of Dunnet with 95% of confidence intervals, using GraphPad Prism Software. P-value < 0.05 was considered statistically significant.

## Results

### ~ 50% of water reservoirs in the Brazilian northeast had cyanobacteria and saxitoxin

Data describing the incidence of cyanobacteria in water reservoirs in Brazil were organized by the percentage of measurements per municipality in the concentration’s ranges: below 10,000 cells/mL, between 10,000 and 20,000 cells/mL and above 20,000 cells/mL. Between 2014 and 2018, NE showed ~ 34% of the measurements above 20,000 cells/mL, while other regions showed no more than 10% of the measurements on this range (Fig 1A – black bar).

**Fig. 1.**
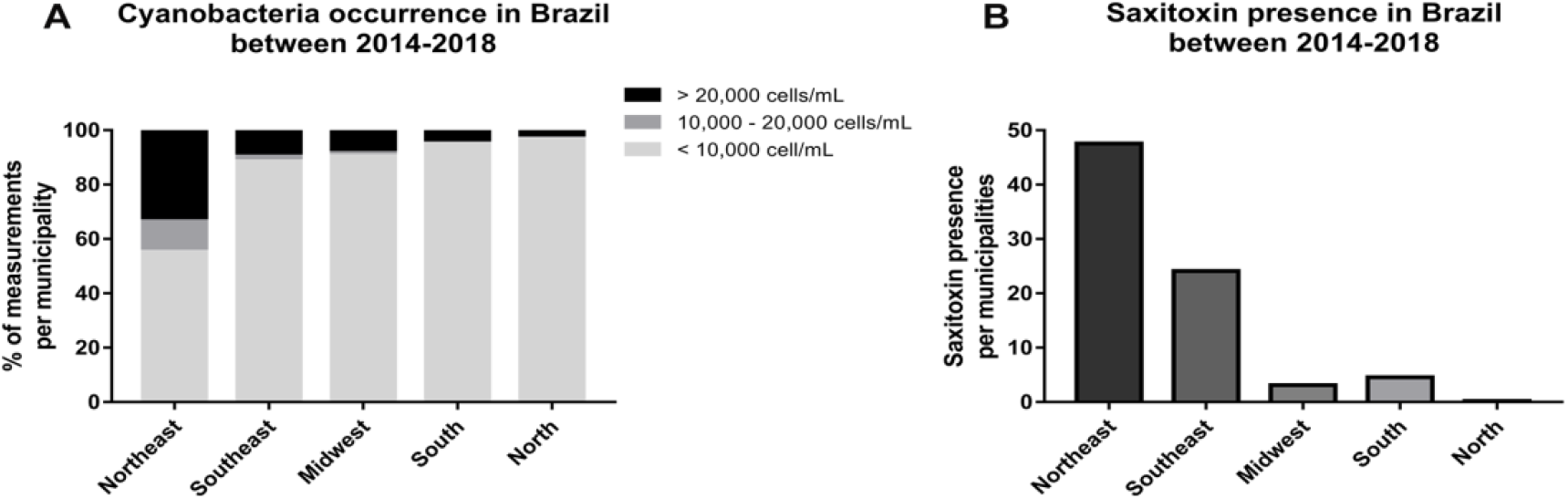
Cyanobacterial and STX occurrence among Brazilian Regions. (A) Cyanobacteria in Brazil between 2014-2018. The measurements per municipality were split in ranges of cyanobacterial concentration. (B) STX 2014-2018 in Brazil. Note that NE had almost twice saxitoxin than SE.

The presence of STX per municipality was also evaluated. Half of NE municipalities had STX in water reservoirs (Fig 1B – dark gray bar), followed by 25% in the SE (Fig 1B – medium gray bar). Other Brazilian regions presented STX in less than 5% of their municipalities (Fig 1B).

### Cell death induced by ZIKV was exacerbated by STX *in vitro* and *in vivo*

In order to evaluate the effects of STX in the live human neural tissue, 50 day-old brain organoids were exposed to 12 μg/L of STX for 13 days and then infected with ZIKV (MOI 0.5, which corresponds to 2−6.5 × 10^5^ plaque-forming unit - PFU - per organoid) (Fig 2A). This concentration of STX was chosen since it was often described in untreated water sources during droughts in the NE of Brazil (32). Fixed organoids were sectioned in cryostat and immunostaining to identify apoptotic cells (TUNEL) and progenitors (Nestin) was performed. ZIKV-infected brain organoids exposed to STX presented ~ 2.5 times more dead cells per mm^2^ than ZIKV-infected organoids (Fig 2B-C). STX alone did not increase cell death in brain organoids (Fig 2C).

**Fig 2.**
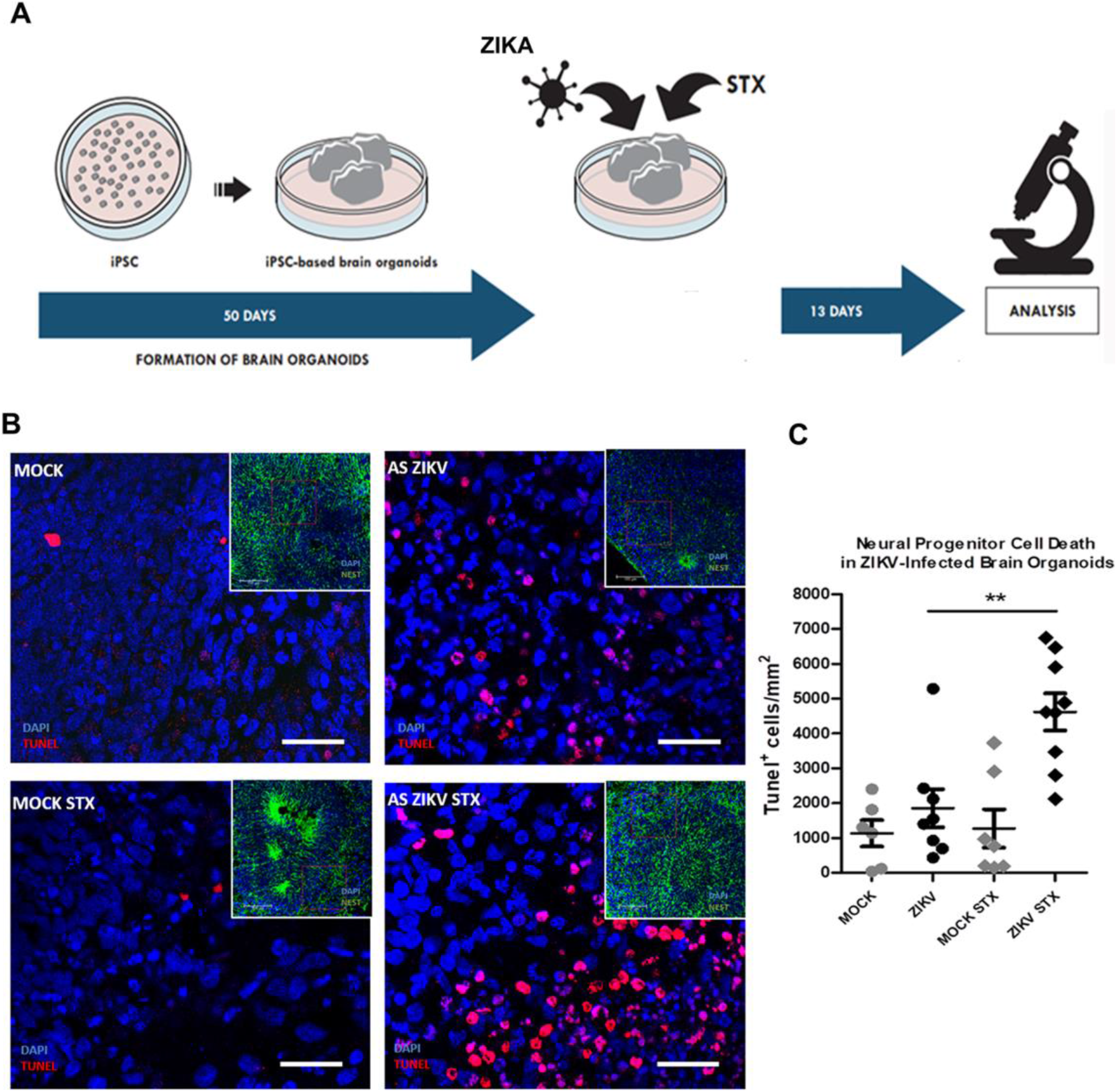
Saxitoxin increases cell death in ZIKV-infected brain organoids. 50-day-old brain organoids were infected with ZIKV and exposed to STX for 13 days. (A) The illustration shows the experimental timeline. (B) Representative images of Nestin-positive areas (green) of untreated or STX-treated Mock and ZIKV-infected organoids. Inserts showing TUNEL-positive cells in areas were used for evaluation. (C) Number of TUNEL-positive cells per Nestin-positive brain organoid areas (mean ± SEM) ** p < 0.01.

To confirm the effects of STX as a cofactor of ZIKV neurotoxicity observed in brain organoids, we used C57BL/6J mice as model. These wild-type animals, due to their efficient type I interferon signaling and ability to control ZIKV replication (30,31), do not present neurological significant impairments derived from vertical ZIKV transmission during embryogenesis (33). Since the population of Brazilian NE is continuously exposed to STX (Fig 1B), and there is insufficient information about their accumulative effect, we decided to analyze the effect of chronic exposition to a low concentration of STX. We offered water contaminated with 15 ng/L of STX to immunocompetent C57BL/6J females one week before mating and continued during gestation. On gestational day 12, females were infected by intraperitoneal injection of 10^6^ PFU per animal. Offspring brains were analyzed on the day of birth (P0) (Fig 3A).

**Fig 3.**
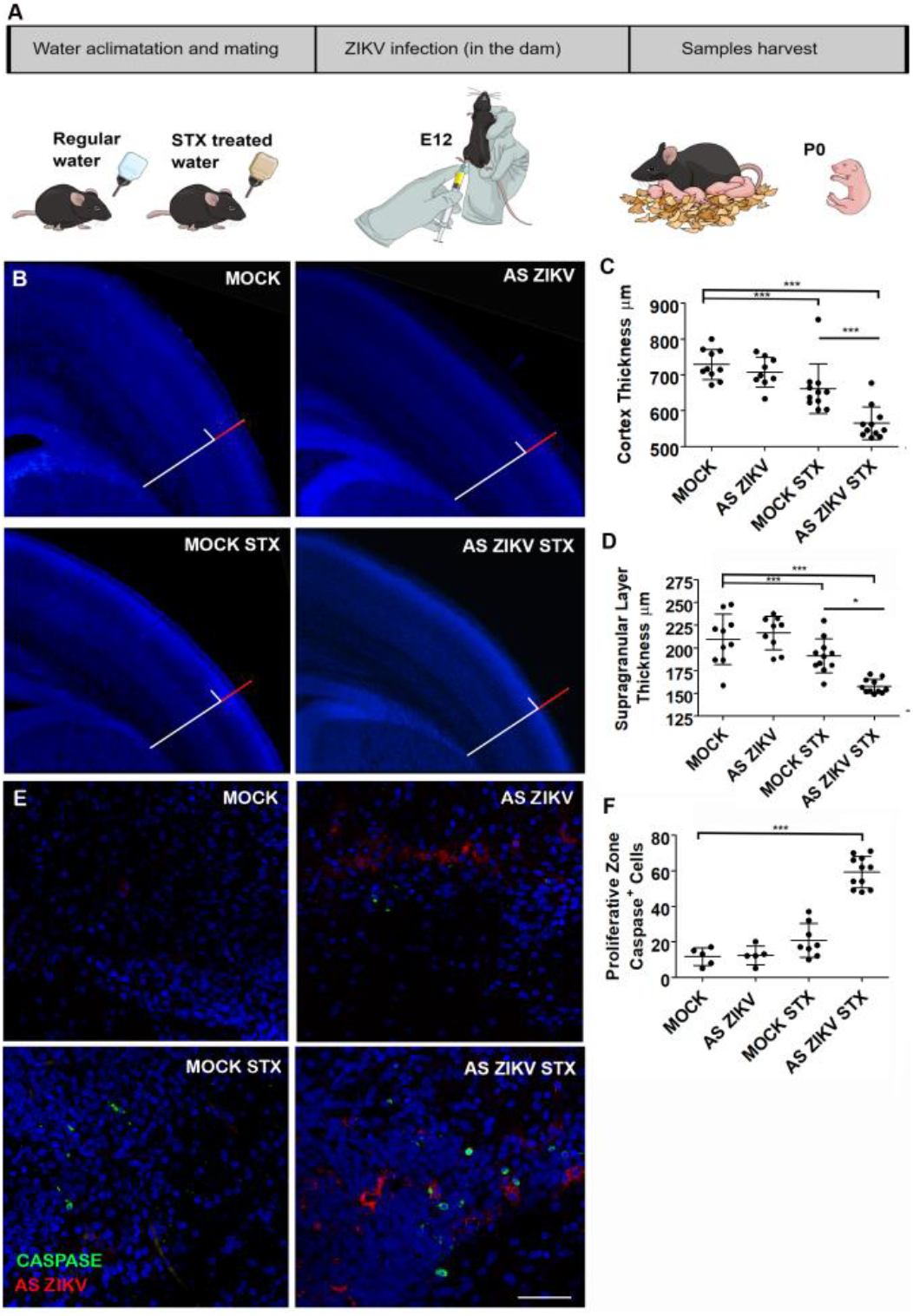
Mice offspring chronically exposed to STX during pregnancy presents severe cortex malformation. C57BL/6J pregnant mice continuously exposed to STX were infected with 10^6^ PFU of ZIKV intraperitoneally at E12. (A) Illustration showing the experimental timeline. (B) Representative somatosensory P0 cerebral cortex sections stained with DAPI. In the Mock group image, the red bar represents the supragranular cortical layers, and the white scale bar represents the remaining cortical layers. The control colored scale bars were replicated in all images for comparison. Cortex (C) and supragranular cortical (D) layers thickness evaluation (mean ± SEM). (E) Representative brain slices stained with cleaved-caspase 3 (green) and NS1 ZIKV (red) in P0 proliferative zones. (F) Quantification of caspase-positive cells in P0 (mean ± SEM) *** p< 0.001, ** p< 0.01 and *p< 0.05.

As expected, the offspring of ZIKV-infected female mice presented mild cortical erosion, while ZIKV infected mice exposed to STX-contaminated water (ZIKV STX group) displayed a significant reduction in cortical thickness, ~ 30% thinner than control animals (Fig 3B, C). Differently than observed *in vitro*, STX alone produced significant effects in cortex thickness (Fig 3C). The thinner cortical layer was predominantly related to a reduction in the supra granular layers (Fig 3D). To confirm if co-exposure of STX and ZIKV induce cell death in the developing cerebral cortex of mice, we quantified the number of caspase-positive cells in their proliferative zones. The amount of cell death in STX ZIKV-infected neonates increased more than twice, in comparison to the other groups (Fig 3E-F).

Our data show that STX exacerbates cell death in the progenitor zones of ZIKV-infected human brain organoids and mice. Therefore, since the incidence of STX in water reservoirs was extremely high in the northeast, and it aggravates the neurogenic impairment caused by ZIKV both *in vitro* and *in vivo*, cyanobacteria may be considered a cofactor to the malformations caused by ZIKV in Brazil.

## Discussion

In the present study, we aimed to determine the possible participation of STX, one of the most neurotoxic and widespread PST naturally found, as a co-insult to ZIKV malformations. First, we showed that cyanobacteria and STX are notably widespread in the Brazilian NE (Fig 1A, B). Moreover, the evaluation of STX and ZIKV association showed a two-fold increase in cell death (Fig 2B, C), while the chronic exposition to a lower concentration of STX in ZIKV-infected pregnant mice revealed a microcephaly-like phenotype.

Issues related to drinking-water contaminated with cyanobacteria have already occurred in Brazil, United States and Australia (3). Toxic cyanobacterial blooms commonly occur in NE of Brazil, where large amount of cyanobacteria and STX are common (Fig 1A, B). A recent study with cyanotoxin-contaminated water from the Brazilian NE showed a deep impairment of zebrafish development, including spine deformation and an increased rate of lethality (34). A previous work showed that neuronal cells exposed to low doses of STX had inhibited axonal-like extensions, suggesting that cells remained in an immature state (15). It has already been shown that neural activity during development prevents inappropriate connections in the brain (35). In human brains, ZIKV infects neural stem cells and glial cells rather than neurons (36). We showed increased cell death in neural progenitors from STX exposed ZIKV-infected brain organoids (Fig 2B, C). It remains to be known the mechanisms by which STX acts in human neural progenitors.

Chronic exposure to STX before and during ZIKV-infection in mice, mimicked what might have occurred in the NE of Brazil. We offered water contaminated with 15 ng/L of STX to pregnant mice. This concentration is considered safe to humans by Brazilian regulatory legislation (16) and is usually found in the drinkable water of the NE of Brazil, according to the SisAgua databank (Ministry of Health). Even in this concentration, significant impairment in cortical thickness (Fig 3B-D) and increased cell death in proliferative zones (Fig 3E, F) were observed in ZIKV-infected mice exposed to STX. STX alone reduced the cortical thickness of mice offspring (Fig 3C) as well, similar to observed in zebrafish (34).

The synergism between cyanobacteria and ZIKV raises awareness that the exposure to STX should also be considered as a public health concern during arbovirus outbreaks. It is important to clarify that microcephaly and other ZIKV-derived congenital abnormalities are multifactorial, therefore, other risk factors may have contributed to foster the uncommon pattern of CZS in Brazil (27). ZIKV outbreaks occurred elsewhere; however, no epidemiological relationship between STX-producing cyanobacteria and congenital malformations derived from ZIKV infection was showed until now.

With this study, we shed light on the importance of governmental regulations for monitoring cyanobacterial blooms and their removal during water treatment, particularly on droughts. We also observed that SXT may act synergistically with ZIKV even at concentrations considered to be safe by Brazilian authorities. Stringent standards and surveillance of drinking water in areas where ZIKV is reported will be critical for minimizing future harmful arbovirus-associated effects on human populations.

## Acknowledgements

This project was supported by Fundação de Amparo à Pesquisa do Estado do Rio de Janeiro (E-26/201.340/2016), Coordenação de Aperfeiçoamento de Pessoal de Nível Superior (88887.116625/2016-01 and 440909/2016-3), Conselho Nacional de Desenvolvimento Científico e Tecnológico (440909/2016-3 and 441096/2016-6), Banco Nacional de Desenvolvimento Econômico e Social (10.2.0051.1), Financiadora de Estudos e Projetos (01.08.0657.00 and 01.12.0161.00), Departamento de Vigilância em Saúde Ambiental eSaúde do Trabalhador – DSAST (25380.001612/2017-70) and intramural grants from D’Or Institute for research and Education. The authors would like to thank Marcia Triunfol and Teresa Puig for helpful discussion during manuscript elaboration; Renan Duarte dos Santos, Camila Vicente Bonfim and Luiz Felipe Lomanto Santa Cruz from Department of Surveillance in Environmental Health and Worker’s Health (DSAST/SVS), from the Brazilian Ministry of Health for providing data of SisAgua databank.

## Supplemental information

**S1 Fig.**
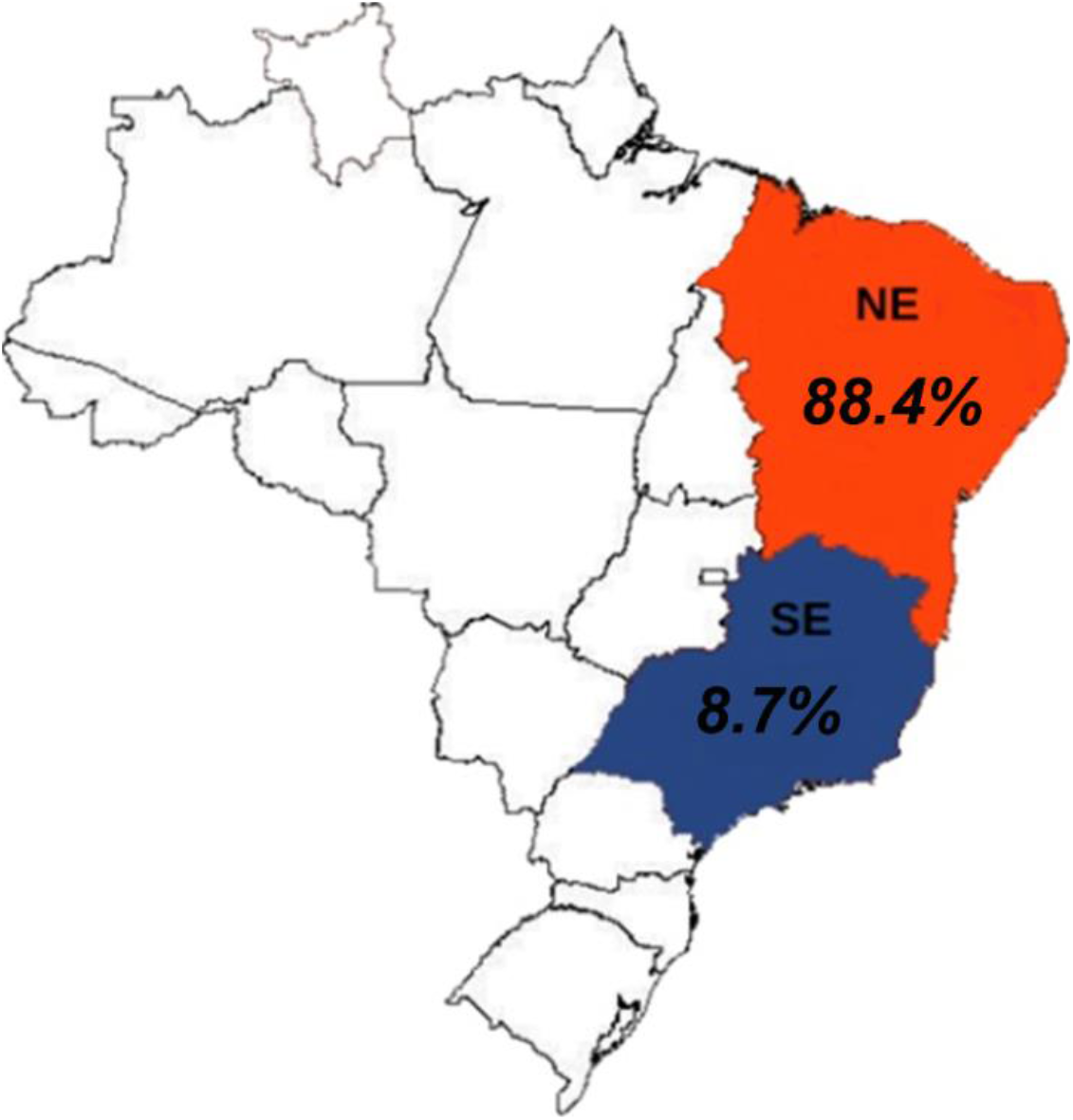
Percentage of brain image exams in the literature reporting microcephaly in NE and SE regions of Brazil. The comparative systematic review selected 37 manuscripts with brain images of infants with ZIKV-related malformations. The percentage of microcephaly brain in exams was placed in a representative map of Brazil, in which southeast region is blue and northeast region is red.

## S1 Method

**Comparative systematic review of ZIKV-derived microcephaly in SE and NE Brazil**

The words “Magnetic Resonance Image”, “Computed tomography”, “Ultrasound” and their acronyms in English and Portuguese, combined with the words “Zika” and “Brazil” were used to search on scientific publication databases PubMed/MEDLINE, LILACS and Scielo. Only manuscripts with SE or NE as the geographical origin of cases (37 manuscripts, including 9 with cases from the SE) were included in analysis. The analysis considered publications until June 2018 in which infants were exposed to ZIKV during the gestational period. The relative percentage of microcephaly-positive brain images per region was obtained using brain exams from each region as 100%.

